# The role of urothelial PIEZO ion channels in urinary function

**DOI:** 10.64898/2026.10.10.758063

**Authors:** Yasmeen M. F. Hamed, Max A. Odem, Shriya Pendyala, Tammy B. Kwok, Kara L. Marshall

## Abstract

The urothelium is increasingly recognized as a dynamic sensory interface that contributes to the regulation of urinary tract function. Mechanosensitive PIEZO ion channels expressed by urothelial cells have been proposed to play important roles in bladder fullness sensation and urinary reflexes. However, the functional contribution of urothelial PIEZO channels to these processes remains unclear. Here, we investigated the consequences of urothelial *Piezo1* and *Piezo2* deletion using the commonly used *Upk2-cre* transgenic line. Surprisingly, combined deletion of urothelial PIEZO channels did not alter urinary behavior, and mice lacking urothelial PIEZO1/2 exhibited normal bladder pressure-volume relationships, indicating preserved bladder filling and voiding function. Notably, *Upk2-cre*; *Piezo1*^*fl/fl*^ mice displayed enhanced external urethral sphincter reflex activity during voiding, suggesting that loss of PIEZO1 from cells targeted by *Upk2-cre* may influence sensory signaling outside of the bladder urothelium. Consistent with this possibility, we identified substantial off-target *Cre* expression in dorsal root ganglia. These findings demonstrate that the *Upk2-cre* line is not restricted to the urothelium and therefore raises important considerations for interpreting associated phenotypes attributed specifically to urothelial gene deletion. Biologically, this raises the possibility that PIEZO1 does not influence bladder pressure-volume handling but might contribute to urinary reflexes.

**HIGHLIGHTS:**

- Urothelial *Piezo1/2* deletion using the *Upk2-cre* line does not alter urinary behavior
- Mice lacking urothelial PIEZO channels show normal bladder pressure-volume handling
- Voiding urethral reflexes are enhanced in *Upk2-cre*^*+*^; *Piezo1*^*fl/fl*^ mice
- *Upk2-cre* line shows off-target *cre* expression in dorsal root ganglia

## INTRODUCTION

Normal urinary bladder function relies on sensing increasing luminal pressure/volume during filling and relaying this information to the central nervous system to coordinate voiding. This process requires precise coordination between the bladder itself and the external urethral sphincter (EUS), mediated by distinct afferent and efferent pathways in the pelvic and pudendal nerves^1,2^. The urothelium, which is the epithelium lining the bladder lumen, has emerged as more than a passive barrier where it is increasingly recognized as a specialized mechanosensory tissue that detects stretch during filling, transducing this signal via afferent nerve fibers^3,4^. Among the candidate mechanotransducers expressed by urothelial cells, the PIEZO family of mechanically activated cation channels, PIEZO1 and PIEZO2^5^, has attracted particular attention^6-8^, given their well-established roles as bona fide mechanosensors in other tissues^9^.

A critical role for PIEZO2 in urination was previously defined^6-8^. Deletion of sensory neuron *Piezo2* in mice impairs low-threshold bladder-stretch sensing and disrupts the micturition reflex, and humans with loss-of-function *PIEZO2* mutations report deficient bladder-filling sensation and impaired bladder control^8^. Subsequent work using *Upk2-cre* to specifically delete *Piezo1* and *Piezo2* in urothelial cells reported subtly altered voiding behavior in double knockout animals, which was only observed during the active dark cycle, reinforcing the idea that PIEZO channels within the urothelium contribute to bladder fullness sensation and function^7^.

The functional role of PIEZO1 in the urothelium remains poorly defined despite its abundant expression throughout all layers of the urothelium and other bladder tissues^6,8^. *In vitro*, PIEZO1 has been proposed to act upstream of stretch-evoked Ca^2+^ influx and ATP release in cultured urothelial cells, positioning it as a candidate mechanosensor linking bladder stretch to purinergic signaling^4^. However, *in vivo* evidence for this role has been inconsistent: urothelial-specific deletion of *Piezo1* has been reported to produce only a limited phenotype, with native urothelial ATP release and voiding behavior largely preserved, raising the possibility that PIEZO1 is dispensable, or that its contribution is masked by compensatory mechanisms^7^. This is at odds with pharmacological evidence that direct activation of PIEZO1 with the agonist Yoda1 alters voiding frequency *in vivo*, suggesting that gain- and loss-of-function approaches to PIEZO1 have yielded discrepant conclusions about its physiological necessity^10^. More recently, urothelial PIEZO1 has also been implicated in pathological bladder remodeling, where its activation was reported to drive inflammation, epithelial-mesenchymal transition, and fibrosis following bladder outlet obstruction^11^, further underscoring that PIEZO1’s contribution to normal, as opposed to pathological, bladder physiology remains unclear. In parallel, we have previously reported that PIEZO1 is also expressed in bladder smooth muscle, where it may contribute to detrusor mechanotransduction independent of the urothelium^12^, raising the possibility that PIEZO1 signaling relevant to bladder function may extend beyond urothelial cells.

Thus, whether and how urothelial PIEZO1 contributes to bladder mechanotransduction and urinary reflexes *in vivo* remains unresolved, and its role continues to be debated across studies and disease contexts. Compounding this uncertainty, prior *in vivo* studies of urothelial PIEZO1/2 function have relied exclusively on the *Upk2-cre* driver, used as a urothelial-specific line^13-16^. Interestingly, there is some evidence of expression in certain brain regions in this line, but PIEZO ion channel expression in the brain is limited and not expected to play a role in physiology^13,14,17^.

Given the uncertain and potentially compensatory relationship between urothelial PIEZO1 and PIEZO2, we sought to further investigate the contribution of urothelial PIEZO channels to urinary tract function, examining both individual and combined loss of *Piezo1* and *Piezo2* using the *Upk2-cre* line to evaluate consequences for bladder filling, voiding behavior, and urethral sphincter reflex activity.

## RESULTS

### Loss of PIEZO1 does not alter intravesicular bladder pressure handling

To determine whether urothelial PIEZO1 contributes to the initiation, strength, or frequency of bladder contractions during voiding, we performed *in vivo* anesthetized cystometry in PIEZO1 knockout (*Upk2-cre*^*+*^; *Piezo1*^*fl/fl*^, P1KO) mice and wildtype littermates. We developed a custom analysis code to parse urinary events by defining windows around inflection points rather than peak-finding. Pressure traces revealed comparable patterns of filling-associated bladder contractions between control and P1KO animals, with no obvious differences in the amplitude or frequency of pressure peaks (Figure 1A). Consistent with these observations, analysis of individual micturition cycles showed no significant differences in peak intravesical pressure between control and P1KO mice (Figure 1B, C) or total pressure generated during individual micturition cycles (Figure 1D).

**Figure 1.**
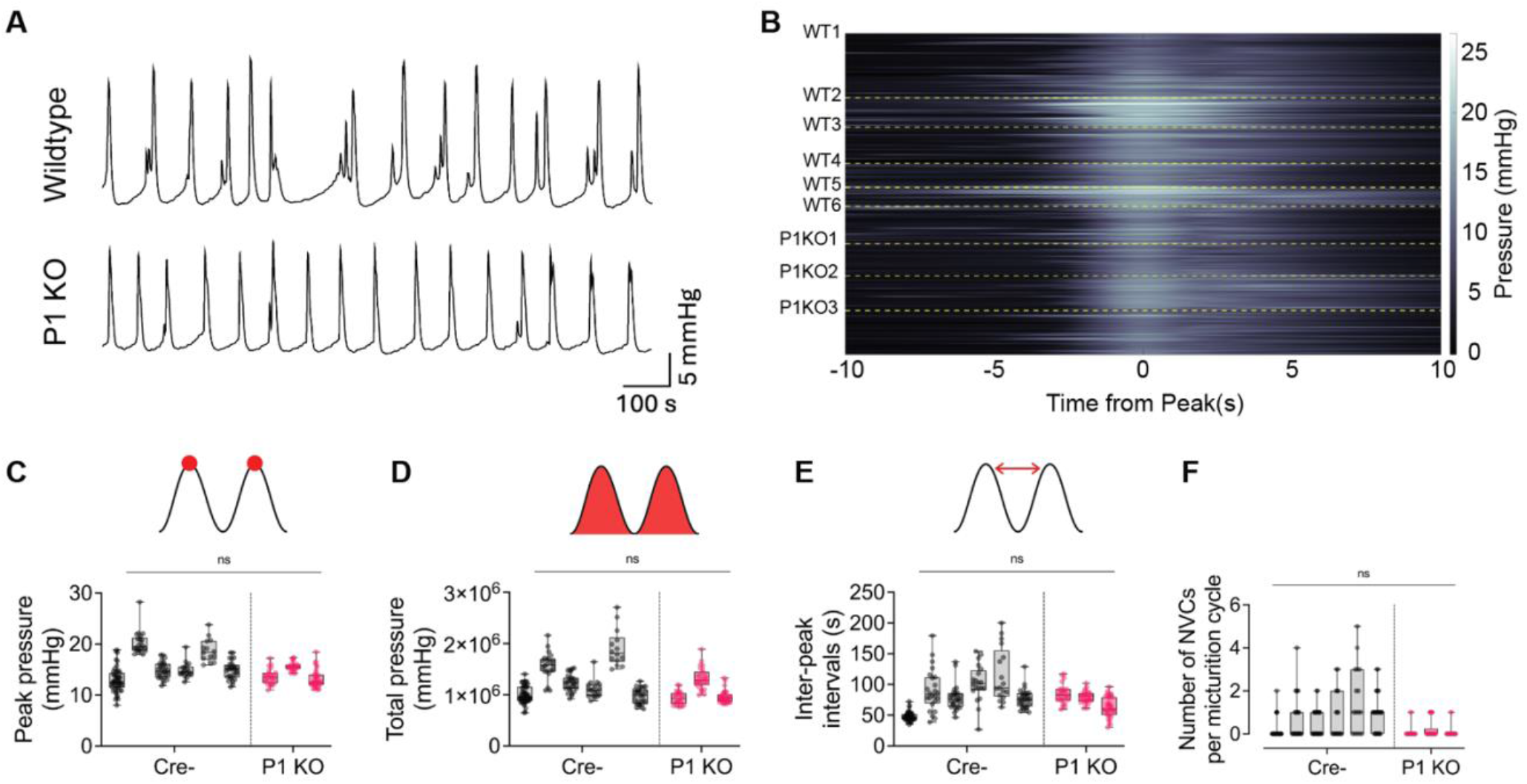
Loss of PIEZO1 does not alter intravesicular bladder pressure handling. (A) Example 20-minute cystometry recordings from a *Upk2-cre*^*–/–*^ control “Wildtype” mouse and a *Upk2-cre*^*+*^; *Piezo1*^*fl/fl*^ knock-out “P1KO” mouse, quantified in C-F. (B) Heatmap showing pressure 10 seconds before and after micturition cycle peaks (scale: black = 0 mmHg, white = 25 mmHg). Each row represents an individual pressure peak, with dashed yellow lines separating individual animals. (C) Peak intravesicular bladder pressure during micturition cycles. (D) Total pressure of individual micturition cycles, calculated as total area under the curve. (E) Inter-peak intervals representing time between pressure peaks of individual mucturition cycles. (F) PIEZO1 KO mice exhibited a trending decrease in the number of non-voiding contractions compared to Cre negative control mice. For all plots: Cre-males *n*=6; P1KO males *n*=3. Box and Whisker plots: box shows 25th to 75th percentiles, middle line shows the median, and whiskers show minimum and maximum values; each box and whisker shows an individual animal, dots show individual pressure peaks. Statistical test: (C-F) Nested *t*-test. (ns) *p* > 0.05, (*) *p* ≤ 0.05, (**) *p* ≤ 0.01, (***) *p* ≤ 0.001, (****) *p* ≤ 0.0001.

We next examined the temporal characteristics of bladder cycles. The interval between successive pressure peaks was not significantly different between control and P1KO mice, indicating that loss of PIEZO1 does not substantially alter the timing of micturition cycles in anesthetized mice (Figure 1E). Finally, we quantified non-voiding contractions (NVCs), which reflect bladder contractions occurring independently of a voiding event. P1KO mice exhibited a trend toward very consistent contractions with a reduced number of NVCs per micturition cycle compared with control animals; however, this difference did not reach statistical significance (Figure 1F). Together, these findings indicate that loss of urothelial PIEZO1 does not significantly alter intravesical pressure generation, contraction dynamics, or micturition cycle timing, suggesting that PIEZO1 is not required for the fundamental filling-sensitive bladder reflex properties during micturition cycles.

### *Piezo1* deletion enhances urethral sphincter activity during bladder voiding

To determine whether urothelial PIEZO1 contributes to coordination between bladder contraction and urethral sphincter activity during voiding, we paired anesthetized *in vivo* cystometry with simultaneous external urethral sphincter (EUS) electromyography (EMG) in control and *Upk2-Cre*^*+*^ P1KO mice. EMG recordings aligned to intravesicular bladder pressure demonstrated robust EUS activity at the initiation of bladder contraction in both genotypes (Figure 2A). Bladder contraction initiation was defined via detection of a peak in the third temporal derivative of pressure. This approach better approximated the pressure inflection point compared to the first and second temporal derivatives. Visualization of EUS activity across individual micturition cycles revealed enhanced overall urethral activity in P1KO mice compared with controls (Figure 2B).

**Figure 2.**
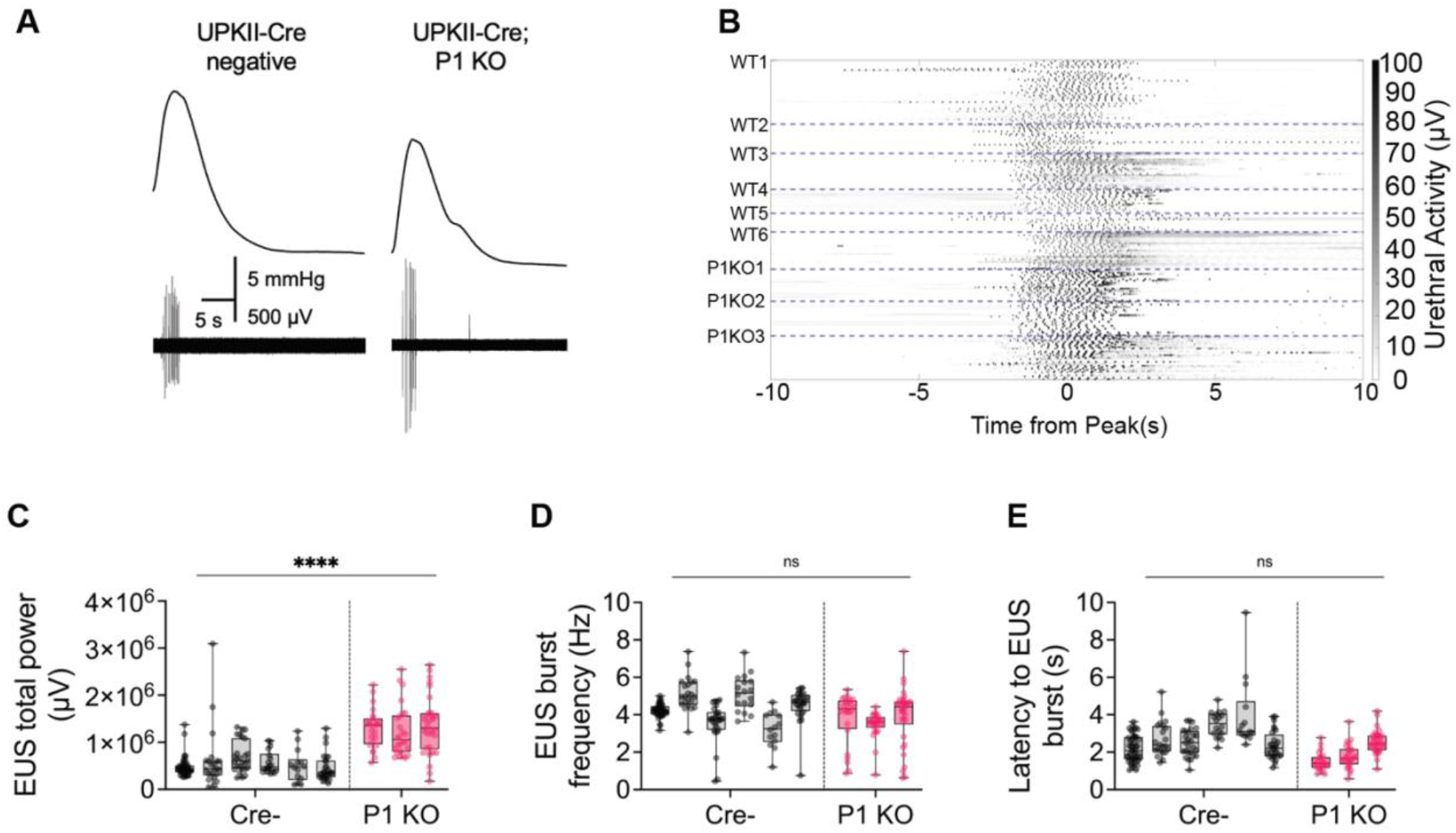
*Piezo1* deletion results in more robust urethral response during bladder voiding. (A) Example external urethral sphincter (EUS) electromyography (EMG) recordings correcponding to pressure peaks during micturition cycles from a *Upk2-cre*^*–/–*^ control “Wildtype” mouse and a *Upk2-cre*^*+*^; *Piezo1*^*fl/fl*^ knock-out “P1KO” mouse. (B) Heatmap showing urethral EMG activity 10 seconds before and after corresponding micturition cycle pressure peaks (scale: white = 0 µV, black = 100 µV). Each row represents EMG activity corresponding to a pressure peak, with dashed blue lines separating individual animals. (C) Total EUS activity during micturition cycles in µV. (D) Frequency of EUS activity bursts during micturition cycles in Hz. (E) Latency time between first EUS activity bursts and corresponding pressure peaks in seconds. For all plots: Cre-males *n*=6; P1KO males *n*=3. Box and Whisker plots: box shows 25th to 75th percentiles, middle line shows the median, and whiskers show minimum and maximum values; each box and whisker shows an individual animal, dots show individual pressure peaks. Statistical test: (C-E) Nested *t*-test. (ns) *p* > 0.05, (*) *p* ≤ 0.05, (**) *p* ≤ 0.01, (***) *p* ≤ 0.001, (****) *p* ≤ 0.0001.

Quantification of total EUS activity during micturition cycles confirmed a significant increase in EUS activity following *Piezo1* deletion (Figure 2C). In contrast, the frequency of EUS activity bursts was not significantly different between control and P1KO mice (Figure 2D), indicating that the increased overall activity was not attributable to an increase in burst frequency. Similarly, the latency between bladder pressure peaks and the onset of the first corresponding EUS activity burst was comparable between genotypes (Figure 2E), as was the pressure threshold required for EUS response and the bladder pressure Tau, defined here as time needed for peak pressure to drop by ∼63.2% (Figure S1).

Together, these findings indicate that deletion of urothelial PIEZO1 using the *Upk2-cre* line enhances the robustness of EUS activity during bladder voiding without significantly altering the firing frequency, threshold, or latency of urethral sphincter bursts. The EUS activity is driven by bladder-evoked stretch reflexes, which are initiated by fast neuronal signals rather than slower urothelial-cell–mediated processes. Thus, these data also point to the possibility that *Upk2-Cre*–mediated targeting of urothelial PIEZO channels might have off-target effects on PIEZO channels in sensory neurons. Hence, we next aimed to probe this possibility.

### *Upk2-cre* mice lacking *Piezo1/2* show intact urinary behavior and preserved bladder pressure-volume handling

Previous studies revealed that *Piezo2* deletion in caudal tissues results in severe urinary behavior deficits in mice^8^, but this was in the context of impaired reflexes. As all reflexes in *Upk2-cre* P1KO were preserved or enhanced, we did not anticipate this would impair urinary behavior or control. In line with this, urinary behavior in *Upk2-cre* P1KO, as well as *Piezo1/Piezo2* double knockout (DKO) animals was comparable to control littermates (Figure S2). Previous work revealed impaired reflexes in *Upk2-cre* P2 KO alone^8^, but the combination of deleting both ion channels together was not sufficient to alter urinary behavior. These observations recapitulate previous findings using the *Upk2-cre* DKO bladders which showed no major changes in urinary behavior, only total volume changes in the dark phase^7^.

One possibility to explain enhanced EUS reflexes in the absence of changes to bladder stretch reflexes is that bladder accommodation to filling is altered, leading to more or less stretch with a given fluid volume. To assess whether urothelium alone can alter pressure-volume (P-V) handling, which is part of what drives reflex sensitivity, we performed an *ex vivo* assay to remove extrinsic innervation and test how urothelial mechanosensation affects P-V handling^12,18^ (Figure 3). Our data indicated no differences in bladder weights, which suggests no major bladder remodeling (Figure 3A). Additionally, we saw no differences in all P-V parameters evaluated, including infused volume to reach 20 mmHg, average compliance, or duration of each phase of the micturition cycle (Figure 3B-F). As we found no bladder-intrinsic changes in pressure handling or bladder-specific reflex alterations, this opens up the possibility that the alterations we observe in the EUS are occurring due to reflex circuitry downstream of the bladder.

**Figure 3.**
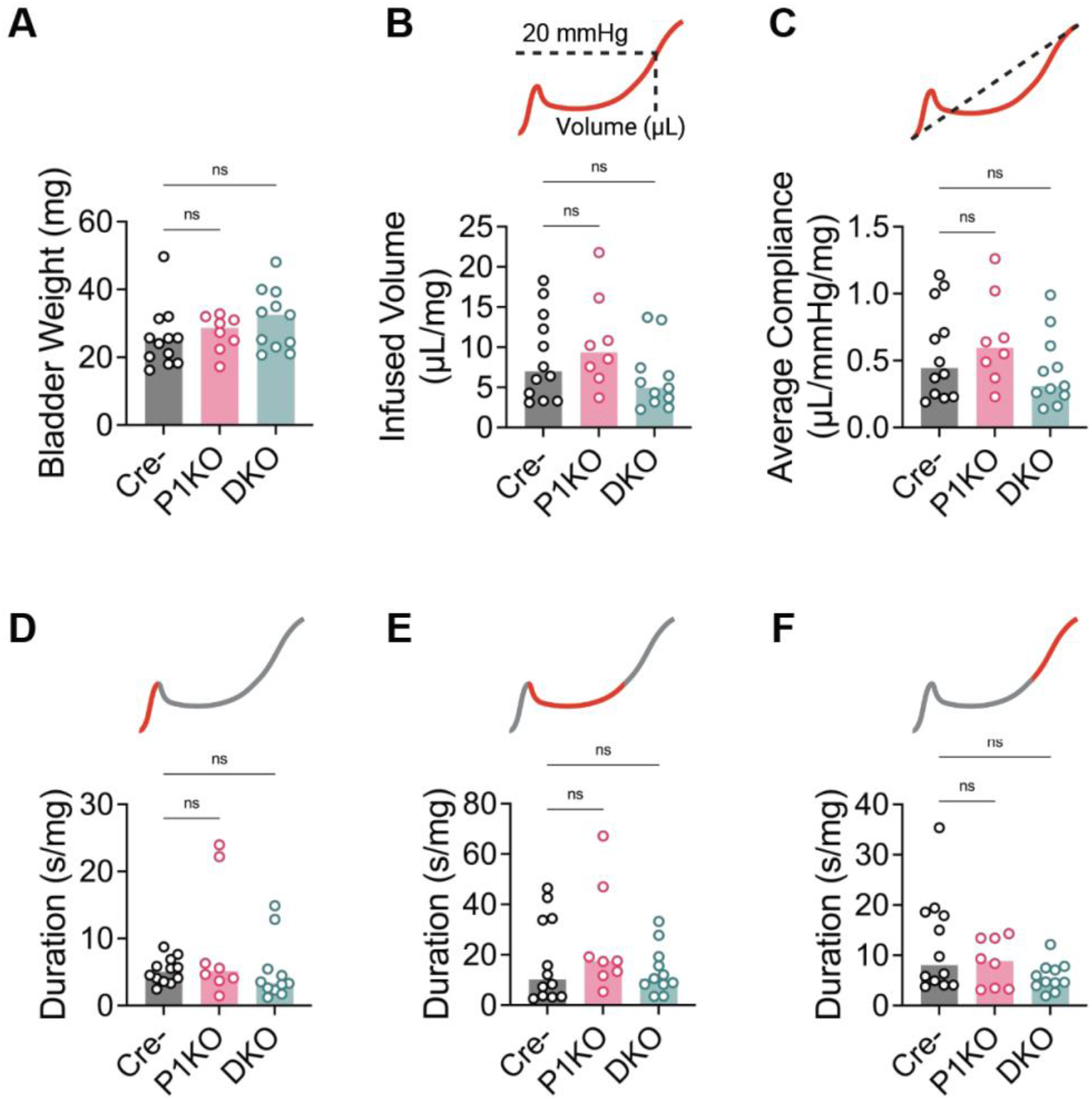
*Upk2-cre* mediated *Piezo1/2* deletion mice show preserved bladder pressure-volume handling in *ex vivo* assay. (A) Bladder weights of male and female mice; *Upk2-cre*^*–/–*^ controls “Cre-” (gray), *Upk2-cre*^*+*^; *Piezo1*^*fl/fl*^ knock-outs “P1KO” (pink), and *Upk2-cre*^*+*^; *Piezo1*^*fl/fl*^; *Piezo2*^*fl/fl*^ double knock-outs “DKO” (green); *n*=8-12/group. (B) Infused volume needed for bladders to reach voiding pressure point (20 mmHg), normalized to empty bladder weight. (C) Average bladder compliance, calculated as ΔV/ΔP over the 0–20 mmHg filling range of pressure– volume (P–V) curves, normalized to empty bladder weight. (D-F) Duration of each bladder cycle phase, normalized to empty bladder weight. For all bar plots: bars show mean; dots show individual animals. Statistical test: (A, D, F) Kruskal-Wallis test with Dunn’s multiple comparisons ad hoc test, (B, C, E) Brown-Forsythe and Welch ANOVA tests with Dunnett T3 correction for multiple comparisons. (ns) *p* > 0.05, (*) *p* ≤ 0.05, (**) *p* ≤ 0.01, (***) *p* ≤ 0.001, (****) *p* ≤ 0.0001.

### *Upk2-cre* drives off-target recombination in dorsal root ganglia and peripheral nerve fibers innervating the bladder

The *Upk2-cre* transgenic line is widely used to achieve urothelium-specific gene deletion, based on the broadly adopted perspective that *Upk2* expression is restricted to the umbrella cells of the urothelium^13-15^. To directly test the specificity of this line, we crossed *Upk2-cre* mice with the *Ai9* tdTomato Cre-reporter line and examined recombination in the bladder and dorsal root ganglia. As expected, *Upk2-cr*^*+*^; *Ai9* bladders showed robust tdTomato labeling throughout the urothelium, confirming efficient on-target recombination in this tissue, while no tdTomato signal was detected in urothelium from *Upk2-cre*^*−*^; *Ai9* littermate controls (Figure 4A).

**Figure 4.**
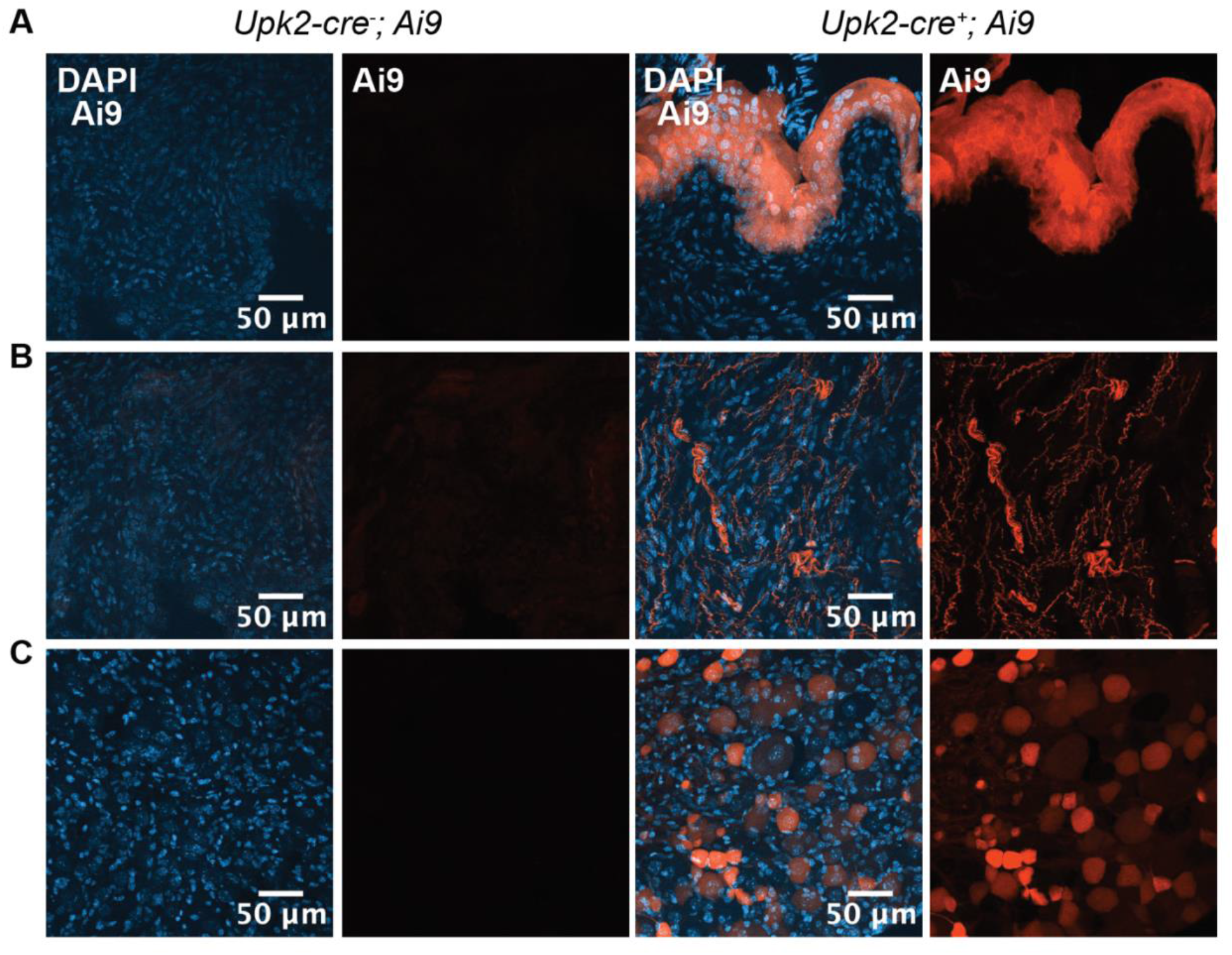
Cre reporter line reveals *Upk2-cre*^*+*^ animals might have off target expresion of cre recombinase in dorsal root ganglia. (A-C) Representative images from *Upk2-cre*^*–*^ (control, left) and *Upk2-cre*^*+*^ (right) mice crossed with the cre reporter line, Ai9. Td-tomato (red) signal indicating Cre recombinase activity is seen in on-target cells, the bladder urothelium (A), as well as off-target cells, including neuronal fibers in the bladder detrussor muscle (B) and dorsal root ganglia (C). DAPI staining shown in blue. Scale bar is 50 µm. *n*=4/group.

Surprisingly, with enhanced exposure, we also observed prominent tdTomato labeling in neuronal fibers within the bladder detrusor muscle of *Upk2-cre*^*+*^; *Ai9* mice. Sensory neurons do not derive from *Upk2*-expressing cells nor do they express *Upk2* after development^19^ (Figure 4B). Given that these labeled fibers likely represent afferent or efferent nerve processes originating from cell bodies outside the bladder wall, we next examined dorsal root ganglia (DRG), the location of bladder-innervating sensory neuron cell bodies. We found that tdTomato^+^ neurons were readily detected in DRG from *Upk2-cre*^*+*^; *Ai9* mice (Figure 4C), strongly suggesting that *Upk2-cre* drives recombination in sensory neurons outside the urothelium.

To confirm that this off-target reporter activity reflects genuine Cre expression rather than leaky or developmental recombination of the Ai9 allele, we performed fluorescent *in situ* hybridization for *Cre* and *Piezo2* transcripts in DRG from *Upk2-cre*^*−*^; *Piezo2*^*fl/fl*^ and *Upk2-cre*^*+*^; *Piezo2*^*fl/fl*^ mice. *Piezo2* transcript was detected in DRG neurons of both genotypes, consistent with the wide expression of *Piezo2* in somatosensory neurons (Figure 5A, B). *Cre* transcript, however, was undetectable in DRG neurons from *Upk2-cre*^*−*^ controls (Figure 5A) but was clearly present in a subset of DRG neurons from *Upk2-cre*^*+*^ mice (Figure 5B). Nonetheless, we did not observe a significant change in FM1-43 signal in DRGs tissues (Figure S3), which labels PIEZO2 activity in sensory neurons^20^. Together, these findings demonstrate that the *Upk2-cre* line is not restricted to the bladder urothelium but also drives recombination in a subset of DRG sensory neurons.

**Figure 5.**
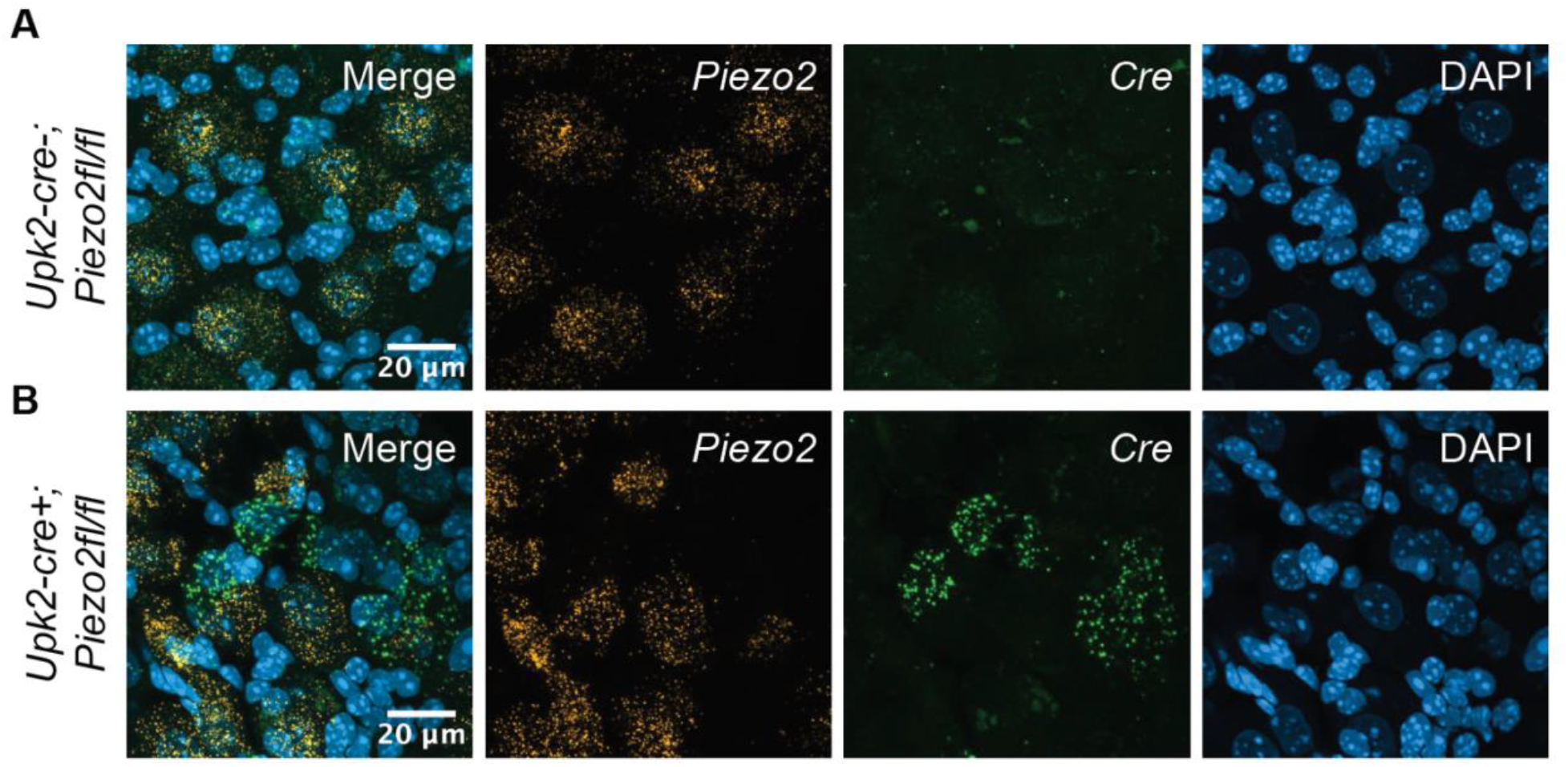
Fluorescent in situ hybridization confirms the expression of cre recombinase in dorsal root ganglia of *Upk2-cre*^*+*^ animals. (A-B) Representative images from fluorescent *in situ* hybridization showing the expression of *Peizo2* (yellow) and *Cre* (green) transcripts in dorsal root ganlgia from *Upk2-cre*^*–/–*^ controls (A) and *Upk2-cre*^*+*^; *Piezo2*^*fl/fl*^ knock-outs (B). DAPI staining shown in blue. Scale bar is 20 µm. *n*=2/group.

## DISCUSSION

The urothelium is a multifaceted epithelial layer, serving as a protective barrier, a dynamic tissue allowing the bladder to expand multiple folds its size, and a sensory tissue that can detect bladder distension and communicate this information to surrounding cells and sensory pathways^3^. The mechanosensitive ion channels, PIEZO2 and PIEZO2, are particularly well positioned to play a key role in urothelial mechanosensory function because of their ability of convert stretch into biological signals^4,7^. PIEZO2 has also been implicated in bladder fullness sensation and urinary reflexes via its established role in bladder-innervating sensory neurons^8^, while both urothelial PIEZO channels have been proposed to contribute to urinary control^7,8^. Our findings suggest that urothelial PIEZO1 and PIEZO2 should be viewed as components of a broader bladder sensory signaling mechanism rather than obligatory determinants of urinary function. Deletion of PIEZO1, or both PIEZO1 and PIEZO2, from the urothelium produced little detectable consequences across the functional measures examined here. This distinction is important because mechanosensitive signaling within urothelial cells^4,7^ does not necessarily imply that these channels are required for the systems-level downstream processes governing bladder pressure-volume handling (*in vivo* or *ex vivo*), voiding behavior, or urinary reflexes.

Our results and interpretation differ from previous studies in which *Upk2-Cre*–mediated deletion of urothelial PIEZO2, alone^8^ or combined with PIEZO1^7^, results in bladder pressure handling, urinary behavior and reflex abnormalities. These divergent outcomes can in part be explained by differing experimental conditions (e.g., duration of urinary behavior assay), biological determinants (e.g., sex, age), and physiological contexts (e.g., circadian state during test). In addition, an important consideration is the specificity of the *Upk2-Cre* line. Our findings indicate that there could be Cre recombinase activity in dorsal root ganglia (DRG) sensory neurons, which warrants caution when interpreting phenotypes resulting from the *Upk2-Cre*–mediated deletion of urothelial genes which are also functional in sensory neurons. This finding is particularly relevant to PIEZO2, given its established and broad functional roles in DRGs^9^ and urinary phenotypes previously reported using *Upk2-Cre*^*7,8*^. Although our FM1-43 measurements do not indicate a generalized disruption of PIEZO2 activity in DRGs, they do not exclude functional recombination of *Piezo2* in a subset of sensory neurons. Thus, taken together, our data suggests that these phenotypes could be, at least in part, mediated by sensory neuronal, rather than or in addition to, urothelial PIEZO2.

The same consideration applies to our observation that enhanced external urethral sphincter (EUS) activity following *Piezo1* deletion. PIEZO1 is expressed and functional in a subset of DRG neurons^21,22^, providing a plausible neural substrate through which off-target recombination could mediate this phenotype. Nonetheless, whether bladder-innervating sensory neurons are among the affected population, and whether neuronal PIEZO1 contributed to the EUS phenotype, remain unresolved. Thus, the phenotype cannot presently be assigned (exclusively) to urothelial PIEZO1. More selective, neuron-sparing genetic approaches will be needed to distinguish urothelial signaling from neuronal mechanosensation, and to define which components of urinary sensory and reflex control genuinely depend on urothelial PIEZO channels.

## Supplementary Figures

**Figure S1.**
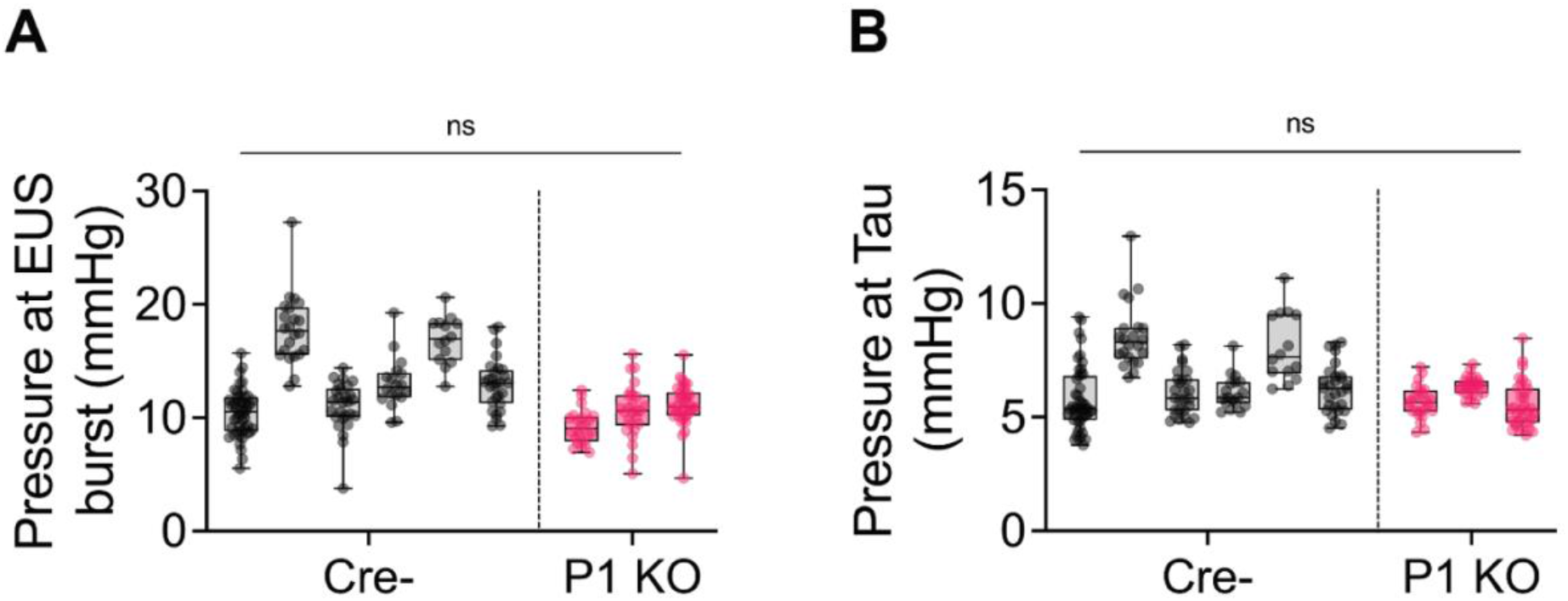
Cystometry with urethral EMG shows no change in coordinated bladder-urethra response parameters in *Upk2-cre; Piezo1*^*fl/fl*^ mice. (A) Pressure at the onset of external urethral sphincter (EUS) burst corresponding to voiding pressure peaks. (B) Pressure at Tau, defined as the time needed for pressure peak to drop by 63.2%. For all plots: Cre-males *n*=6; P1KO males *n*=3. Box and Whisker plots: box shows 25th to 75th percentiles, middle line shows the median, and whiskers show minimum and maximum values; each box and whisker shows an individual animal, dots show individual pressure peaks. Statistical test: (A-B) Nested *t*-test. (ns) *p* > 0.05, (*) *p* ≤ 0.05, (**) *p* ≤ 0.01, (***) *p* ≤ 0.001, (****) *p* ≤ 0.0001.

**Figure S2.**
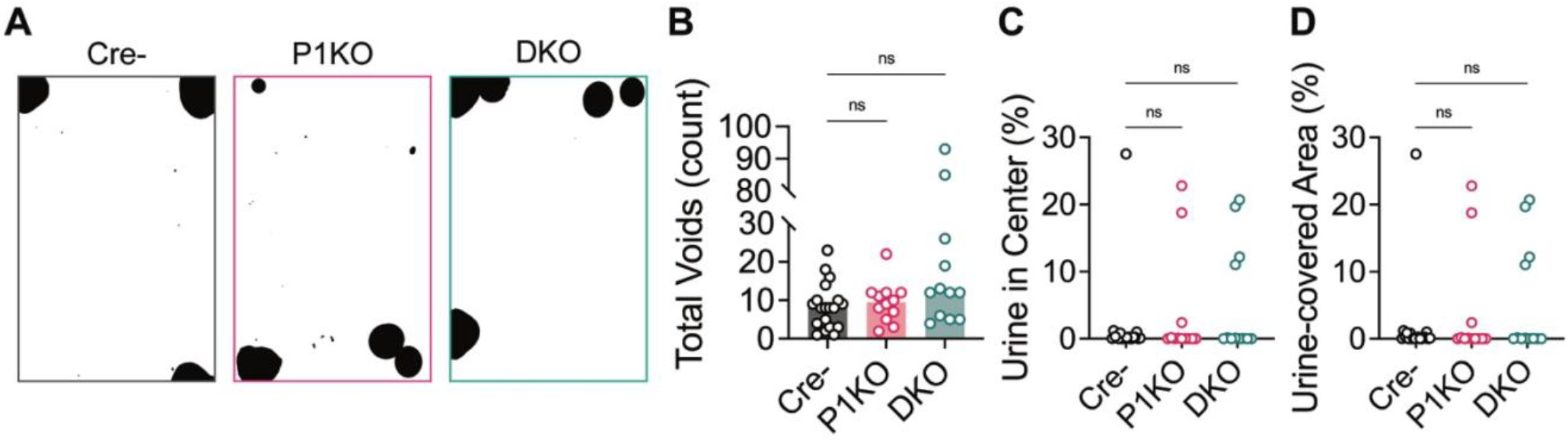
*Upk2*-mediated *Piezo1/2* knock out does not alter urinary behavior. (A) Representative end-point images of 4 h void spot assay papers from male and female mice; *Upk2-cre*^*–/–*^ controls “Cre-” (left, gray frame), *Upk2-cre*^*+*^; *Piezo1*^*fl/fl*^ knock-outs “P1KO” (center, pink frame), and *Upk2-cre*^*+*^; *Piezo1*^*fl/fl*^; *Piezo2*^*fl/fl*^ double knock-outs “DKO” (right, green frame); imaged under UV light. Urine void spots are shown in black; quantified in B-D (*n*=12-19/group). (B) Total number of urinary void spots deposited in a 4 h assay. (C) Percentage of urine deposited in center 25% of cage area in a 4 h void spot assay. (D) Total urine-covered area at the end of a 4 h void spot assay, quantified as percentage of total cage area. For all bar plots: bars show mean; dots show individual animals. (B-D) Statistical test: Kruskal-Wallis test with Dunn’s multiple comparisons ad hoc test. (ns) *p* > 0.05, (*) *p* ≤ 0.05, (**) *p* ≤ 0.01, (***) *p* ≤ 0.001, (****) *p* ≤ 0.0001.

**Figure S3.**
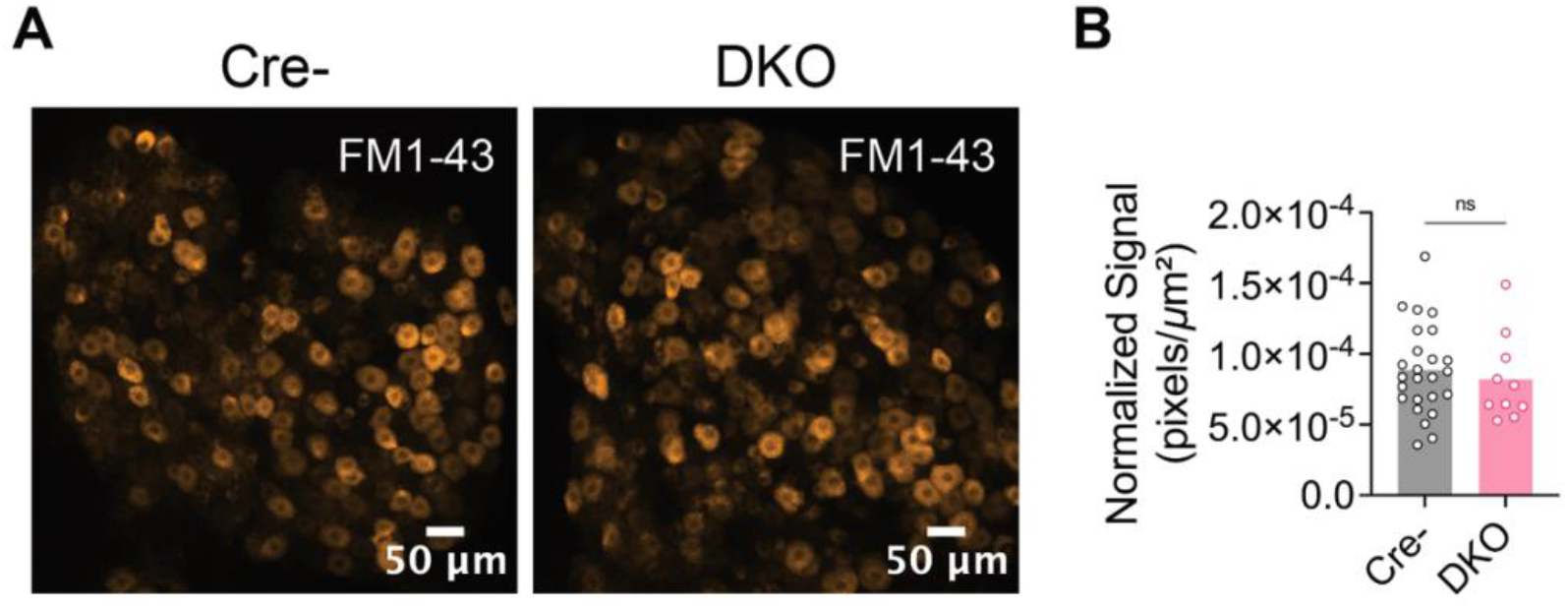
FM1-43 labelling does not indicate significnant changes in overall PIEZO2 activity within *Upk2-cre*^*+*^; DKO dorsal root ganglia sensory neurons. (A)Representative images showing PIEZO2 activity in mouse dorsal root ganglia (DRGs) tissues from *Upk2-cre*^*–/–*^ controls “Cre-” and *Upk2-cre*^*+*^; *Piezo1*^*fl/fl*^; *Piezo2*^*fl/fl*^ double knock-outs “DKO”, labelled with the fluorescent styryl dye FM1-43. (B)FM1-43 signal intensity in Cre- and DKO mouse DRGs, normalize to tissue area. Bars show mean; dots show individual DRG images. Five images were analyzed per animal. *n*=2-3 animals/group. (B) Statistical test: Welch’s *t*-test. (ns) *p* > 0.05, (*) *p* ≤ 0.05, (**) *p* ≤ 0.01, (***) *p* ≤ 0.001, (****) *p* ≤ 0.0001.

## METHODS

### Experimental Models

#### Mice

For all experiments apart from the *in vivo* cytometry, mice were housed in Baylor College of Medicine’s clean Transgenic Mouse Facility (TMF) in accordance with the approved IACUC guidelines (14 h/10 h light/dark cycle at 21 ºC with 40-70% humidity). Mice were always given ad libitum access to food and water. For *in vivo* cytometry experiments, mice were housed in the animal facility at Scripps Research (La Jolla, CA) under standard housing conditions (12:12h light:dark cycle, 22ºC room temperature, 30-80% humidity (not controlled), and given ad libitum access to food and water. To obtain urothelial *Piezo1* knockout (P1KO) or *Piezo1/Piezo2* double knock-out (DKO) mice, *B6(129)-Tg(Upk2-cre)1Rkl/WghJ* animals (The Jackson Laboratories, stain #029281)^15^ were crossed with *B6*.*Cg-Piezo1*^*tm2*.*1Apat*^*/J* line (The Jackson Laboratories, strain #029213)^23^, with or without *B6(SJL)-Piezo2*^*tm2*.*2Apat*^*/J* (The Jackson Laboratories, strain #027720)^24^. To obtain mice carrying a Cre reporter, *B6(129)-Tg(Upk2-cre)1Rkl/WghJ* animals (The Jackson Laboratories, stain #029281)^15^ animals were crossed to Ai95 *B6;129S-Gt(ROSA)26Sor*^*tm95*.*1(CAG-GCaMP6f)Hze*^*/J* (The Jackson Laboratories, strain #024105).

### Behavioral and Physiological Assays

#### Void spot assay (VSA)

Urinary behavior studies were performed as previously described ^12^. Adult male and female mice were placed in custom-made bottomless cages that mimic the animals’ home cage environment (11.5” x 7”), set on thick chromatography paper (05-714-5, Fisher Scientific) in a darkened room and allowed to behave freely. Before testing, mice were acclimated to the full testing set-up and behavior room for 2-3 days, 1h each day. On the test day, mice were allowed to behave freely in the test chambers for 4h without access to food or water to avoid accidental water damage to the test papers or biasing where the animals spend time/choose to urinate. Representative animals from each group were placed on a custom-built table with clear acrylic base which is equipped with video cameras at above and below each cage and UV lights placed underneath the cages to allow video recording for movement tracking analysis and urine-spot visualization in real time. At the end of the test, chromatography papers with urine were imaged while illuminated by UV light. Images were thresholded and converted to B&W binary images using ImageJ (v2.16.0/1.54p), and the total number of black pixels was counted. A region of interest corresponding to the center 25% of cage area was drawn and used to quantify the percent of urine signal (pixels) in the center cage arena. For automated void spot count and classification into primary and secondary voids, IMARIS was used (Imaris x64, v11.0.0, Oxford Instruments).

### Tissue functional assays

#### Ex vivo bladder pressure-volume analysis

A mouse ex vivo bladder prep was set-up, as previously described^23^. Briefly, we utilized the gastrointestinal motility set up from Med Associates Inc. A small organ bath with tubing kit (Med Associates Inc., Cat. #GIM-101 and TBU-GIMM-SUPERFUSE) was continuously perfused with circulating Krebs–bicarbonate solution (KBS, Thermo Scientific, Cat. #J67591.K2) solution (maintained at 37ºC, oxygenated with 95% O_2_/5% CO_2_ gas mix) using a gas dispersion tube (Med Associates Inc., Cat. #GD-L). Continuous circulation was achieved using a digital MasterFlex® L/S® Digital Miniflex® single-channel peristaltic pump (Avantor, Cat. #07525-40). Adult male and female mice were euthanized in accordance with the lab’s approved IACUC protocols, perfused with ice-cold 1X PBS, then the bladders with intact ureters and urethra were immediately collected in pre-oxygenated KBS solution in a black Pyrex bottom Sylgard dissection dish where they were cleaned from attached connective tissues and emptied of urine by gently pressing on the bladder in the apex to urethra direction on a Kimwipe. When possible, both ureters were then tied off using a 6-0 silk suture (Roboz Surgical Instrument Co., Cat. #SUT-14-1). Bladders were transferred to the organ bath. Bladders were then catheterized with a blunt 25G, 0,5” needle (SAI Infusion Technologies, Cat. #B25-50) connected to a single-channel syringe pump (Braintree Scientific Inc., BS-8000), filling the bladder with oxygenated KBS at 15 µL/min. Syringe pump tube was also connected through a three-way stopcock to a pressure sensor (BIOPAC Systems Inc., Cat. #RX104A), which records pressure through a pressure transducer and general purpose transducer amplifier system (BIOPAC Systems Inc., Cat. #TSD104A and DA100C). This set-up allows live recording of intravesicular bladder pressure in response to controlled volume filling using a data acquisition system with AcqKnowledge software (BIOPAC Systems Inc., Cat. #MP160WSW). Finally, pressure-volume curves were analyzed using a custom-made MATLAB code by an experimenter blinded to the animals’ information. Trials with clearly leaking bladders, identified by a sudden significant drop in pressure, were excluded.

#### Cystometry and urethral electromyography (EMG)

Anesthetized *in vivo* bladder cystometry with EMG was performed as previously described in Marshall *et al*., 2020^8^. Male mice were anesthetized with isoflurane dynamic dosing (5% for induction, then gradually decreasing to 1-1.3% for maintenance) using the SomnoSuite system (Kent Scientific). The bladder was catheterized with a saline line connected to a peristaltic syringe pump (continuously pumping at 20 µL/min) and a pressure sensor (Biopac, Cat. #RX104A-MR) which is connected to an MP160 system amplifier (Biopac, Cat. #DA100C) via a pressure transducer (Biopac, Cat. #TSD104A). Tungsten electromyography electrodes were inserted directly into the external urethral muscle (A-M systems, Cat. #795500) and connected to a differential amplifier (Biopac, Cat. #EMG100C), with the following settings: 1000 gain, 10 Hz sample rate, low-pass filter 5 kHz, 60 Hz notch filter, and 100 Hz high-pass filter. Data were only collected once the mouse started stably cycling (bladder filling and voiding), which was roughly 30-40 minutes from the start of recording. Once data collection was about to commence, bladder filling rate was increased from 20 µL/min to 30 µL/min. Data were collected using the Acqknowledge software (v.4.4.2) and processed using custom scripts written in MATLAB (v.2019B/2025B).

### Histology

#### Ai9 Cre reporter activity tracing

Adult *Upk2-cre*^*+*^; *Ai9*^*+*^ and *Upk2-cre*^*–*^; *Ai9*^*+*^ male and female mice were euthanized according to the lab’s IACUC-approved protocols, perfused with ice-cold 20-25 mL 1X phosphate buffered saline (PBS) followed by 20-25 mL ice-cold 4% paraformaldehyde (PFA, diluted in 1X PBS). Bladders were dissected, emptied of urine, then briefly washed in 1X ice-cold PBS before being drop-fixed in 4% PFA overnight at 4ºC (Thermo Scientific, Cat. #047377.9M). Tissues were moved from 4% PFA to 30% sucrose in 1X PBS (Sigma-Aldrich, Cat. #S0389-500G) for cryopreservation and incubated overnight at 4ºC. Bladders were then embedded in Tissue-Tek® O.C.T. Compound (Sakura Finetek, Cat. #4583), and flash frozen in liquid nitrogen. Fixed tissues were then sectioned at 20 µm and collected onto Superfrost™ Plus Microscope Slides (Fisher Scientific, Cat. #12-550-15). Slides were washed with 1X PBS for 5 min. on rocker to remove O.C.T., then stained with DAPI for 10-15 seconds. Finally, tissue slides were mounted using ProLong™ Gold Antifade Mountant (ThermoFisher Scientific, Cat. #P10144) and imaged under a 20X objective on a Zeiss LSM 900 confocal microscope.

### RNAscope

Adult wildtype C57BL/6J mice were euthanized using isoflurane overdose followed by thoracotomy and perfusion with 20 mL ice-cold 1X PBS, in accordance with the IACUC-approved protocol. Dorsal root ganglia (DRG) tissues were harvested into ice-cold 1X PBS before being embedded in Tissue-Tek® O.C.T. Compound (Sakura Finetek, Cat. #4583), and flash frozen in liquid nitrogen. Fresh-frozen tissues were then sectioned at 20 µm and collected onto Superfrost™ Plus Microscope Slides (Fisher Scientific, Cat. #12-550-15). Target transcripts were detected using RNAscope™ Multiplex Fluorescent Reagent Kit v2 (ACD Biotechne, Cat. #323100) following the manufacturer’s protocol for fresh-frozen tissues. The following probes were used: Mm-Piezo2-E43-E45 (ACD Biotechne, Cat. #439971) and Mm-Cre (ACD Biotechne, Cat. #312281). Mouse positive control probe (ACD Biotechne, Cat. #320881) and negative control probe (ACD Biotechne, Cat. #320871) were also used. Signal was detection using TSA Vivid™ dyes (TORCIS Bioscience, Cat. #6456 [Fluoresceine Tyramide] and #6457 [Cyanine 3 Tyramide] diluted in RNAscope™ TSA Buffer (ACD Biotechne, Cat. #322810). Finally, tissue slides were mounted using ProLong™ Gold Antifade Mountant (ThermoFisher Scientific, # P10144) and imaged under a 63X objective on a Zeiss LSM 900 confocal microscope. Tissues from each animal were tested against all probes, and at least 5 images were captured per animal.

### FM styryl dye labeling in dorsal root ganglia

PIEZO2 activity in dorsal root ganglia (DRGs) tissues was visualized using FM styryl dye, as previously described^20^. Briefly, wildtype adult C57BL/6J female mice were injected intraperitoneally with FM1-43FX styryl dye (Invitrogen, Cat. #F35355) prepared at a dose of 1.12 mg/Kg body weight, diluted in 1X sterile PBS. Mice were returned to their normal home cage environment and, approximately 48h later, mice were euthanized and perfused with 1X PBS followed by 4% paraformaldehyde (PFA). DRG tissues were dissected into ice-cold 4% PFA and fixed for at least 1h at 4ºC. Tissues were then whole-mounted onto glass slides in mounting medium (Fluoromount-G™, Invitrogen, Cat. #00-4958-02) and imaged with a 10x objective (0.3 NA) on a confocal microscope (Zeiss, LSM 800).

## ACKNOWLEDGEMENTS

This work was supported by the Howard Hughes Medical Institute’s Freeman Hrabowski Scholars Program (K.L.M.), NIH grant numbers R00DK128621 (K.L.M.) and R01DK142807-01 (K.L.M.), McNair Medical Foundation (K.L.M.), Pew Charitable Trusts (K.L.M.), and Rita Allen Foundation (K.L.M.). The authors would like to thank Dr. Ardem Patapoutian for his generous support while generating the cystometry data.

## Data Statement

All original data will be made available upon request. The cystometry analysis code is deposited on GitHub at https://github.com/KLM-PI/cystometry_matlabscripts

## Author contributions (CRediT)

Y.M.F.H. and K.L.M. conceptualized the study. Y.M.F.H., S.P., T.B.K, and K.L.M. performed experiments. Y.M.F.H., M.A.O., and K.L.M analyzed the data. Y.M.F., M.A.O., and K.L.M. developed the methodology for this work. Y.M.F.H. visualized and validated the data. Y.M.F.H. and K.L.M. supervised the work and wrote the original manuscript draft. Y.M.F.H., M.A.O., and K.L.M. provided edits on the manuscript draft. All authors reviewed the manuscript.

